# Eukaryotic plankton community stability across reef environments in Bocas del Toro Archipelago (Panamá)

**DOI:** 10.1101/750356

**Authors:** Andrea M. Rodas, Logan K. Buie, Hannah E. Aichelman, Karl D. Castillo, Rachel M. Wright, Sarah W. Davies

**Affiliations:** Biology Department, Boston University, Boston, MA; Deparment of Marine Sciences, University of North Carolina at Chapel Hill, Chapel Hill, NC; Environment, Ecology and Energy Program, University of North Carolina at Chapel Hill, Chapel Hill, NC; Department of Genetics, Harvard Medical School, Boston, MA; Department of Biological Sciences, Smith College, Northampton, MA

**Keywords:** coral reefs, plankton, diel vertical migration, phytoplankton, zooplankton, reef zone, metabarcoding, 18S

## Abstract

Variation in light and temperature can influence the genetic diversity and structure of marine plankton communities. While open ocean plankton communities receive much scientific attention, little is known about how environmental variation affects tropical coral reef plankton communities. Here, we characterize eukaryotic plankton communities on coral reefs across the Bocas del Toro Archipelago in Panamá. Temperature loggers were deployed for one year and mid-day light levels were measured to quantify environmental differences across reef zones at four inner and four outer reef sites: Inner: Punta Donato, Smithsonian Tropical Research Institute (STRI) Point, Cristobal, Punta Laurel and Outer: Drago Mar, Bastimentos North, Bastimentos South, and Popa Island. Triplicate vertical plankton tows were collected mid-day and high-throughput 18S ribosomal DNA metabarcoding was leveraged to investigate the relationship between eukaryotic plankton community structure and reef zones. Plankton communities from STRI Point were additionally characterized in the morning (∼08:00), mid-day (∼12:00), and evening (∼16:00) to quantify diel variation within a single site. We found that inshore reefs experienced higher average seawater temperatures, while offshore sites offered higher light levels, presumably associated with reduced water turbidity on reefs further from shore. However, these significant reef zone-specific environmental differences did not correlate with overall plankton community differences or changes in plankton genetic diversity. Instead, we found that time of day within a site and diel vertical migration played structuring roles within these plankton communities, and therefore conclude that the time of community sampling is an important consideration for future studies. Overall, plankton communities in the Bocas del Toro Archipelago appear relatively well mixed across space; however, follow-up studies focusing on more intensive sampling efforts across space and time coupled with techniques that can detect more subtle genetic differences between and within communities will more fully capture plankton dynamics in this region.

## INTRODUCTION

The diversity and abundance of marine plankton communities are well known to be affected by environmental variation including but not limited to temperature, nutrients, and light (Andersson et al. 1994; D’Croz et al. 2005). While open ocean and coastal plankton communities are relatively well-studied, plankton communities inhabiting oligotrophic tropical coral reefs have received far less attention, even though these reefs experience environmental variations that are likely to structure these communities across space and time. For example, organisms inhabiting different reef zones experience strongly divergent environmental conditions (Varela et al. 2001; Castillo et al. 2011; Siegel et al. 2013). Inshore reef zones generally experience greater environmental variation associated with changing tidal cycles, increased mean temperatures driven more restricted flow and shallower reef extent, and reduced salinities and increased turbidity associated with freshwater input from rivers and runoff (Lirman and Fong 2007). Offshore reefs are buffered by the open ocean and thus exhibit clearer seawater with more stable temperatures, resulting in enhanced light penetration generally favoring photosynthetic organisms (Boyer et al. 2015). These physical differences in water quality parameters might be therefore expected to influence the structure of plankton communities on coral reefs. First, because there are species-specific thermal optima for plankton survival (Mauchline 1998), temperature differences across reef zones may play a strong role in structuring plankton communities. Additionally, as light levels are a major factor affecting phytoplankton growth (Harrison and Turpin 1982; Edwards et al. 2016), spatial variation in light availability across reefs can have cascading food web effects that influence the entire ecosystem (Andersson et al. 1994; Barrera-Oro 2002).

Shifts in the structure of plankton communities are considered to be robust bioindicators of subtle environmental changes because species that comprise these communities have rapid life-cycles that allow for quick responses to environmental perturbations (Hays et al. 2005; Richardson 2008). For example, shifts in plankton distributions associated with warming waters were documented in the northeast Atlantic from 1959–2000 (Lindley and Daykin 2005). Furthermore, storms and upwelling events affect local water chemistry by introducing nutrient runoff from land, which can rapidly change the distribution of plankton, ultimately impacting their behavior and growth (Dunstall et al. 1990; Richmond and Woodin 1996). Plankton are also fundamental to a healthy food web, as they provide energy to higher trophic-level organisms such as marine birds, fish and corals (Fenchel 1988; Frederiksen et al. 2006). On coral reefs specifically, plankton are an important source of heterotrophic nutrition to corals. Heterotrophy has been shown to increase coral survival and recovery after heat stress (Johannes et al. 1970; Ferrier-Pagès et al. 2010; Hughes and Grottoli 2013; Tremblay et al. 2016) and to mitigate temperature-induced coral bleaching (Grottoli et al. 2006; Aichelman et al. 2016). However, few studies have examined how these important tropical coral reef plankton communities are influenced by environmental variation across different reef zones (Chiba et al. 2018).

Plankton community surveys began in the early 1800s when the first net suitable for sampling zooplankton was developed (Fraser 1968). Historically, plankton communities were characterized by microscopic examination of each microorganism (Johannes et al. 1970; Irigoien et al. 2004). This method relies on advanced taxonomic abilities of the observer to identify diverse species across different life stages as well as extensive time investment. Other methods for assessing plankton communities include measuring zooplankton organic biomass, which can offer insights into the overall abundance, but not the diversity or taxa-specific abundance, of the sampled community (D’Croz et al. 2005). Recent technological advancements in next-generation sequencing have provided a robust and reliable method to identify and characterize the diversity and relative taxa abundances of plankton communities through high-throughput single locus metabarcoding sequencing (Albaina et al. 2016; Bucklin et al. 2016; Abad et al. 2016). In this approach, a genomic locus homologous across all Eukaryota is amplified and sequenced and unique taxa are identified as amplicon sequence variants (ASVs) based on some threshold of similarity in DNA sequence (Eiler et al. 2013; Lindeque et al. 2013; Kermarrec et al. 2014). This high-throughput analytical method provides information about species presence or absence and relative abundance, based on the number of observed reads mapping to any particular taxa, without the need for morphological examination. The precision of ASV identification by next-generation sequencing continuously improves as the databases used to identify species grow (Quast et al. 2013).

Here, we characterized temperature and light environments of eight reef sites on inshore and offshore reef zones across the Bocas del Toro Archipelago in Panamá. We then leveraged 18S ribosomal DNA metabarcoding to gain quantitative insights into how these environmental conditions influence plankton communities across] these two different reef zones. In addition, we also assessed plankton communities at three timepoints (morning, mid-day, and late afternoon) at a single site to explore diel variations in plankton communities. Overall, these data help capture how heterotrophic opportunities on coral reefs might vary across space and time in this region, which can ultimately affect food webs dynamics in these marine environments.

## MATERIALS AND METHODS

### Abiotic Environmental Conditions in Bocas del Toro

To assess environmental differences across reef zones, we characterized thermal and light profiles at four inshore sites (Punta Donato, Smithsonian Tropical Research Institute [STRI] Point, Cristobal, and Punta Laurel) and four offshore sites (Drago Mar, Bastimentos North, Bastimentos South, and Popa Island) within the Bocas del Toro Archipelago reef complex in Panamá (Fig. 1A). These eight sites were categorized as either inshore or offshore based on their distance from mainland Panamá (Fig. 1A; Table 1). Temperature conditions *in situ* were quantified by deploying data loggers (HOBO Pendant, Onset Computer Corporation) at each sampling site for approximately one year, and temperature data was recorded every fifteen minutes at each site for the duration of the deployment. Logger deployment began at the end of May (STRI Point and Popa Island) to early June (Punta Donato, Cristobal, and Drago Mar) in 2015 and loggers were retrieved in August 2016. Loggers from Cristobal, Punta Donato, STRI Point, Drago Mar, and Popa Island were retrieved, but loggers from Punta Laurel, Bastimentos North, and Bastimentos South were unable to be found and presumed lost. Temperature maximum, minimum, and daily range were averaged over the deployment period for each site where loggers were retrieved (Table 2). Mean daily maximum temperature between June 15–30 were used as a proxy for conditions experienced during plankton community sampling.

**Table 1:** Plankton sampling sites on the Bocas Del Toro Archipelago, Panamá including reef zone, latitude, longitude and date of collection.

| Site | Zone | Lat(°N) | Long(°W) | Date |
| --- | --- | --- | --- | --- |
| Punta Donato | Inshore | 9.41815 | 82.34412 | 6/5/15 |
| STRI Point | Inshore | 9.35198 | 82.26627 | 5/27/15<br>6/3/15<br>6/4/15 |
| Cristobal | Inshore | 9.26237 | 82.2409 | 6/4/15 |
| Punta Laurel | Inshore | 9.13277 | 82.11958 | 6/3/15 |
| Bastimentos North | Offshore | 9.34798 | 82.16798 | 5/29/15 |
| Bastimentos South | Offshore | 9.28747 | 82.09232 | 5/30/15 |
| Popa Island | Offshore | 9.1836 | 82.04942 | 5/31/15 |
| Drago Mar | Offshore | 9.4246 | 82.32468 | 6/1/15 |

**Table 2:**
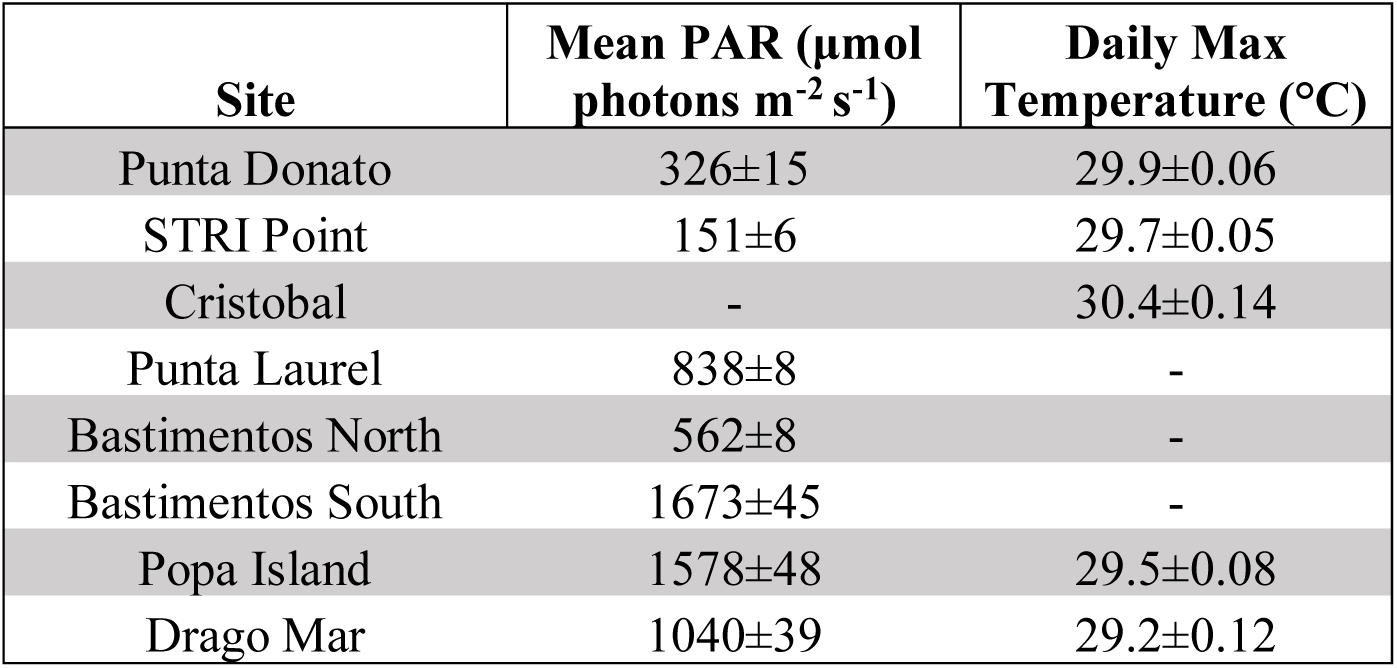
Mean light values (top 20 PAR values) and mean daily temperature maximums +/-SE from 15–30 June 2015 for each reef site where plankton collections were conducted. Cristobal does not have a light value because it was too cloudy on the day of sampling. The HOBO loggers for Punta Laurel, Bastimentos North, and Bastimentos South reef sites were lost during their year of deployment so no temperature data are available for these sites.

| Site | Mean PAR ($\mu\text{mol photons m}^{-2} \text{s}^{-1}$) | Daily Max Temperature ($^{\circ}\text{C}$) |
| --- | --- | --- |
| Punta Donato | $326 \pm 15$ | $29.9 \pm 0.06$ |
| STRI Point | $151 \pm 6$ | $29.7 \pm 0.05$ |
| Cristobal | - | $30.4 \pm 0.14$ |
| Punta Laurel | $838 \pm 8$ | - |
| Bastimentos North | $562 \pm 8$ | - |
| Bastimentos South | $1673 \pm 45$ | - |
| Popa Island | $1578 \pm 48$ | $29.5 \pm 0.08$ |
| Drago Mar | $1040 \pm 39$ | $29.2 \pm 0.12$ |

**Fig. 1.**
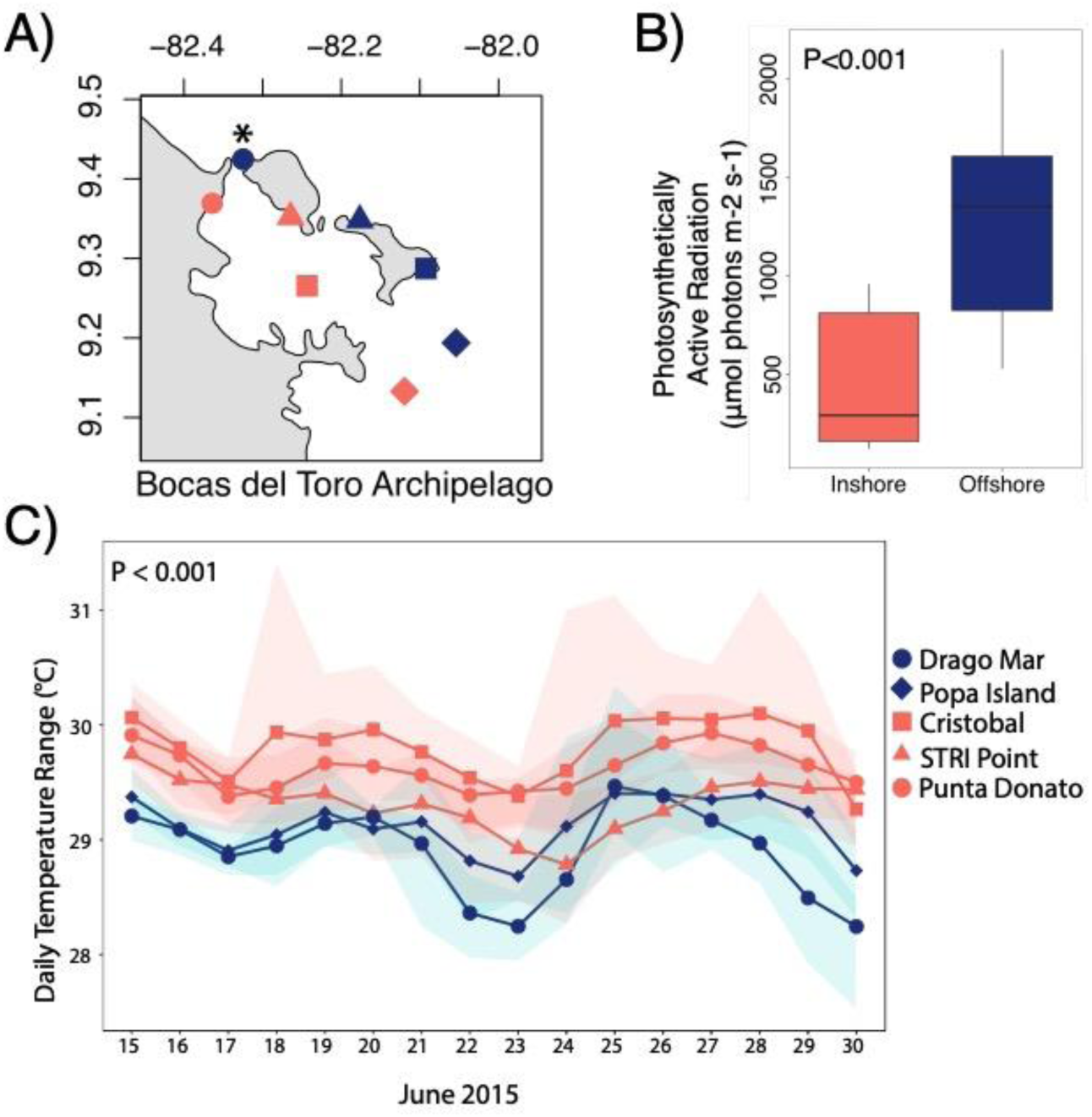
Environmental conditions in Bocas del Toro. (A) Location of collection sites. Symbols indicate site: Punta Donato = salmon circle, STRI Point = salmon triangle, Cristobal = salmon square, Punta Laurel = salmon diamond, Drago Mar = blue circle, Bastimentos North = blue triangle, Bastimentos South = blue square, Popa Island = blue diamond. (B) Mean maximum daily light values averaged across reef zone. Error bars indicate the minimum and maximum light levels. (C) Daily temperature ranges for each site (inshore = salmon, offshore = blue). Symbols represent mean temperatures. Shaded regions encompass the maximum and minimum values for each site. P-value indicates that inshore reef sites are significantly warmer than offshore reef sites. Drago Mar is additionally indicated with a * to correspond with Figure 2B.

An underwater 2π Quantum Sensor (LI-192, LI-COR Inc.) was used to measure photosynthetically active radiation (PAR) for all sites at the time of plankton sample collections with the exception of Cristobal due to consistently overcast conditions. For the remaining sites, PAR levels were measured every thirty seconds between the hours of approximately 10 a.m. and 2 p.m. on sampling days. To account for variations in daily cloud cover, only the maximum twenty PAR measurements collected from each site were used to compare differences across reef types and reef sites (Table 2). A one-way ANOVA (R Development Core Team 2018) was used to test for differences in mean light level and daily maximum temperature across reef zones (Fig. 1C, B).

### Plankton Community Collections and 18S Metabarcoding Preparations

Between 27 May 2015 and 5 June 2015, three replicate vertical plankton tows were conducted at each of the eight sites using a plankton net with 0.5 m diameter and 60 μm mesh filter. Filtered water was then passed through an additional 100 μm filter to concentrate collections and samples were preserved in 200 proof ethanol at a volume of 50 mL. Samples were brought back to the laboratory at the University of North Carolina at Chapel Hill and maintained at −20°C until DNA isolation.

Two replicate DNA isolations were completed for each plankton tow following the extraction method described in Davies et al. (2013). A subset of each well-mixed plankton sample (1.5 mL) was centrifuged to pellet plankton, after which ethanol was decanted. Plankton were then immersed in DNA digest buffer (100 mM NaCl, 10 mM Tris-Cl pH 8.0, 25 mM EDTA pH 8.0, 0.5 % SDS 5 μL Proteinase-K) for 1 hour at 42°C followed by a standard phenol-chloroform extraction procedure. In brief, an equal volume of 25:24:1 buffer-saturated phenol:cholorform:isoamyl alcohol (PCA) was added to the sample, centrifuged, and the resulting aqueous layer was separated. PCA separation was repeated two additional times to further clean the sample and reduce PCR inhibition. DNA was precipitated using 100% ethanol and 3M NaOAc, rinsed with 80% ethanol, and then resuspended in 50 μL milliQ water. DNA concentrations were quantified using a Nanodrop (model ND1000, Thermo Scientific) and all extracts were visualized on 1% agarose gels to assess DNA integrity.

The 18S rRNA region was targeted in each plankton community using original primers from Stoeck et al. (2010), which were then modified for compatibility with Illumina MiSeq. The forward primer sequence was 5’-<u>TCTCGGCGCTC</u>*AGATGTGTATAAGAGACAGNNNN***CCAGC ASCYGCGGTAATTCC**-3’ and the reverse primer sequence was <u>GTCTCGTGGGCTCGG</u>*AGA TGTGTATAAGAGACAGNNNN*ACTTTCGTTCTTGAT-3’ where the text in bold is the 18S target, italics represents linker sequence and underlined text represents Illumina adapter linker sequences, which bind to Illumina adapters during the second PCR (ESM Fig. 1). Each 20 μL polymerase chain reaction (PCR) mixture contained 0.2 mM dNTP mix, 0.5 U *Extaq* polymerase (Takara Biotechnology), 2.0 μL 10X *Extaq* buffer, 100 ng of DNA template, 0.1 μM forward and reverse primer mix, and 12.4 μL milliQ water. PCR amplification was performed using the following profile: 95°C for 5 minutes, followed by 20 cycles of 95°C for 40 seconds, 59°C for 2 minutes, and 72°C for 1 minute, and then an extension period of 7 minutes at 72°C. To avoid PCR biases, samples were cycle checked as per Quigley et al. (2014) to ensure that all samples were amplified to an equivalent intensity when visualized on a 2% agarose gel. Samples that failed to amplify during were diluted 10× with milliQ water, which yielded successful amplification in all cases. PCR products were purified using a GeneJET PCR purification kit (Fermentas Life Sciences). A second PCR was then performed to incorporate unique barcodes and Illumina adapters into each sample for Illumina MiSeq sequencing following Baumann et al. (2017). The PCR thermal profile for this barcode reaction was the same as that described above; however, only four cycles were used. All samples were then visualized together on the same agarose gel and differing volumes of each barcoded sample were pooled based on band intensities. The resulting pooled library was run on a 1.5% agarose gel and the band was excised and soaked in 30 μL milliQ overnight at 4°C. The liquid eluate was sequenced at University of North Carolina at Chapel Hill’s High-Throughput Sequencing Facility using Illumina MiSeq paired-end 300 base pair (bp) sequencing. All raw reads are archived in the National Center for Biotechnology Information (NCBI) Short Read Archive (SRA) under accession number PRJNA507270.

### Plankton Community Analysis

The R statistical environment (R Development Core Team 2018) was used for all data analyses. Scripts for all environmental and sequencing analyses and all environmental data can be accessed at https://github.com/rachelwright8/planktonCommunities. We implemented the *<u>dada2</u>* package to characterize plankton community genetic diversity and structure (Callahan et al. 2016). First, FASTQ files were trimmed for sequence lengths of 250 bp for forward reads and 200 bp for reverse reads based on quality of reads. The first 24 bp from forward reads and 19 bp from reverse reads (representing primer sequence) and all base pairs with quality scores less than or equal to twenty were truncated from all reads. Identical reads were dereplicated, then matching forward and reverse reads were merged. Merged sequences with lengths outside the 365–386 bp range were removed from the analysis as likely products of non-specific primer binding. Chimeric sequences were also removed, resulting in a total of 39% of the original reads remaining (ESM Table 2; Supplemental Files 1 & 2), which were then assigned taxonomy from the Silva database version 123 (https://www.arb-silva.de) using the assignTaxonomy function in *dada2*, with minimum bootstrap confidences of 5 for assigning a taxonomic level. Minimum bootstrap confidences of 50 were also tested and we observed identical results at both taxonomic levels (taxonomic identities can be found in Supplemental File 3). All downstream analyses here are reported on the bootstrap confidence of 5.

The package *phyloseq* was used to create an amplicon sequence variant (ASV) per sample counts table (McMurdie and Holmes 2013). The ASV file was then separated for all mid-day samples and STRI Point time course samples for two separate sets of analyses. The R package *MCMC.OTU* was used to purge rare ASVs that appeared in fewer than 1% of all samples per Green et al. (2014). ASV count data were then log-normalized and principal coordinate analysis (PCoA) was used to compare plankton communities between reef zones, sites, and time of day using the R package *vegan* (Oksanen et al. 2018). The *adonis* function was used to test for differences in plankton communities across these factors. Lastly, Simpson and Shannon diversities for each plankton sample were calculated using *phyloseq* and then these differences in diversity across reef zones, sites and time of day were compared using ANOVA and Tukey’s HSD tests.

### Variation in Specific Plankton Taxa

Differential abundance analyses were performed on ASV counts using DESeq2 (Love et al. 2016). Two negative binomial models were fit to test for differentially abundant ASVs by reef zone and time of day using the models ASV count ∼reef zone and ASV count ∼ time, respectively. Raw ASV counts are available in Supplementary Files 1 (reef zone) and 2 (time of day). Counts were normalized for size factor differences and a pairwise contrast was computed for inshore and offshore reef zones and between all three pairwise comparisons for time of day. An FDR adjusted p < 0.05 (Benjamini and Hochberg 1995) represents significantly different abundances. To visualize these differences, raw counts were rlog normalized and heatmaps with hierarchical clustering of abundance profiles were created with the *pheatmap* package (Kolde 2018). DESeq2 results are available in Supplemental Files 4 (reef zone) and 5 (time of day) and taxonomic assignment results can be found in Supplemental File 3.

## RESULTS

### Divergent Environmental Conditions across Bocas del Toro Reef Zones

Temperature loggers were retrieved from five of eight sites. Maximum daily temperatures over the first two weeks of deployment were significantly higher at inshore sites than offshore sites (p < 0.001; Fig. 1C). Average temperature and standard error for the first two weeks of deployment at inshore sites was 30.02 ± 0.07°C while the offshore sites had an average of 29.37 ± 0.08°C. The top twenty PAR values (μmol photons m^−2^ s^−1^) recorded at each site show that *in situ* light levels at inshore sites were significantly lower (438 ± 38 μmol photons m^−2^ s^−1^) than offshore sites (1213 ± 54 μmol photons m^−2^ s^−1^) (p < 0.001; Fig. 1B). Overall, inshore sites are warmer and experience lower light levels when compared to offshore sites on Bocas del Toro reefs.

### Plankton Communities Do Not Differ Across Reef Zones

Principal coordinate analysis (PCoA) revealed that overall plankton communities were not significantly different (p = 0.256, Fig. 2A) between inshore and offshore reef zones. Also, inshore and offshore reef zones did not differ in mean Shannon’s Index of Diversity (H) (p = 0.178) or mean Simpson’s Index of Diversity (D) (p = 0.19) (Fig. 2B&C). Although overall plankton communities did not differ by reef zone, there were several taxa significantly enriched in either inshore or offshore reef zones (N = 18), with inshore reef sites exhibiting enrichment of 4 taxa compared to offshore sites exhibiting enrichment of 14 taxa (Fig. 2B). In particular, three out of the four enriched taxa in the inshore sites were photosynthetic organisms (highlighted in green text; Fig 2B). It is also worth noting that Drago Mar exhibited taxa abundance profiles more similar to inshore sites (starred samples; Fig. 2B), which is interesting given its relative proximity to shore relative to other offshore sites (Fig. 1A).

**Fig. 2.**
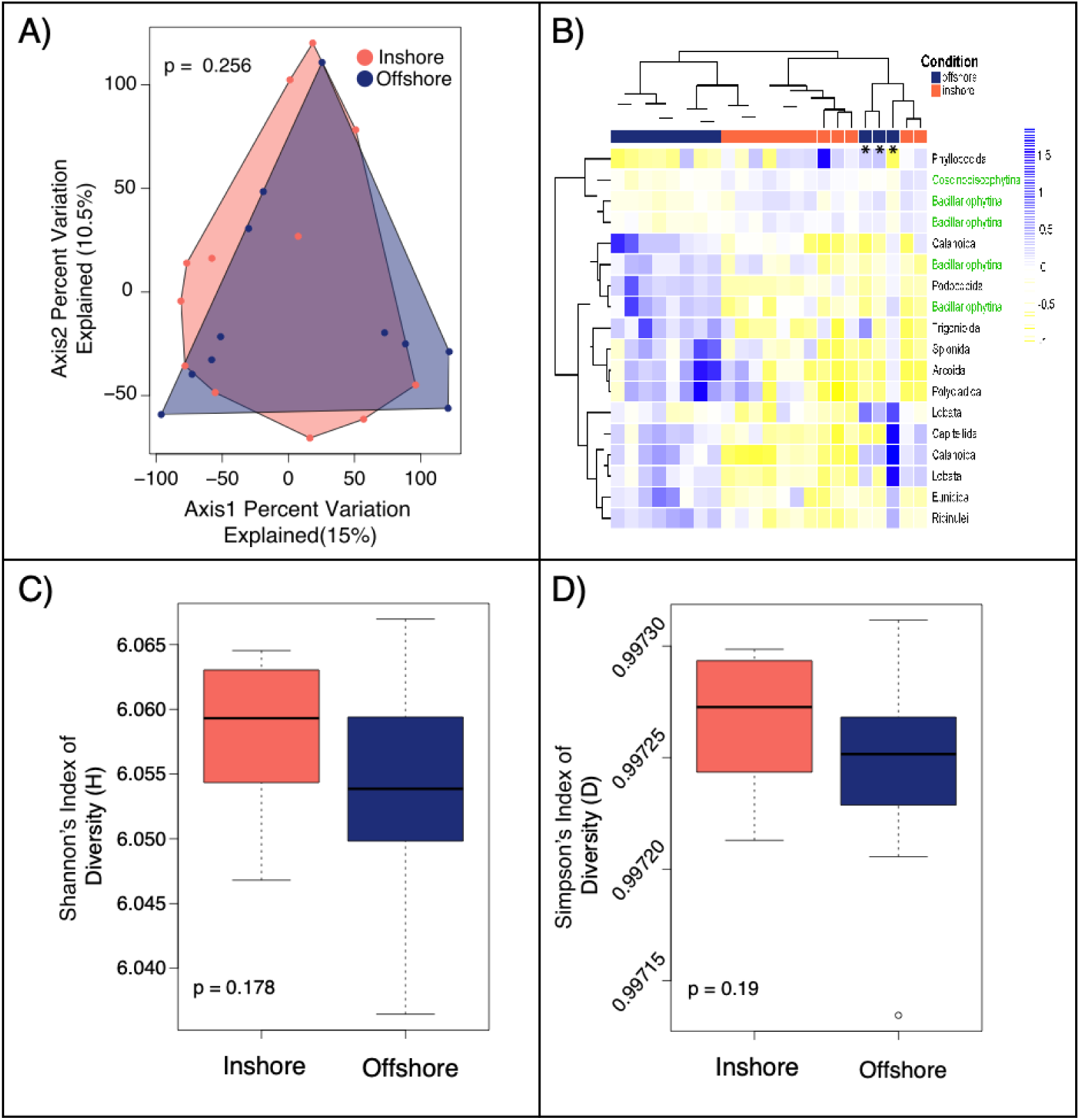
Variation in mid-day plankton samples by reef zone. (A) Principal coordinate analysis of plankton communities by reef zone (inshore/offshore). Percentages on each axis indicate the amount of variation explained by each axis (Inshore = salmon, offshore = blue). P-value indicates results from the *Adonis* test. (B) Heatmap of the most differentially abundant taxa across the two different reef zones. Coral and blue blocks indicate that libraries originated from inshore and offshore plankton communities, respectively. Columns represent unique plankton tows and rows represent differentially abundant taxa. * symbols indicate libraries originating from Drago Mar. Taxa listed in black are heterotrophic whereas taxa listed in green are autotrophic. The color scale is in log2 (blue: enriched, yellow: depleted) and taxa and samples are clustered hierarchically based on Pearson’s correlation of their relative abundance across samples. (C) Mean Shannon and (D) Simpson diversity of plankton communities based on reef zone. P-values demonstrate that there were no statistical differences in diversity across reef zone.

### No Reef Site-specific Differences in Plankton Communities

PCoA grouped by individual reef site indicated that there were no significant differences in plankton communities across individual sites (p = 0.509; ESM Fig. S2A). Furthermore, no significant differences in diversity between sites based on mean Shannon’s Index of Diversity (p = 0.297; ESM Fig. S2B) or mean Simpson’s Index of Diversity (p = 0.385; ESM Fig. S2C) were observed. For diversity indices, means for most sites ranged from 6.05–6.06 for Shannon and 0.99725–0.99727 for Simpson, with the exception of Popa Island, which exhibited the lowest mean diversity for both indices (Shannon: 6.04, Simpson: 0.99720; ESM Fig. S2B, C).

### Time of Day Significantly influenced Plankton Community Composition

PCoA analysis partitioned by time of day revealed that there was a significant shift in plankton community structure across different times of day at STRI Point (early, mid-day, and late; p = 0.003; Fig. 3A). These time course differences were driven by mid-day plankton communities, which were distinct from plankton communities observed at early and late times of day (Fig. 3A). Plankton samples collected mid-day exhibited the least variation in community composition between its three replicate tows (Fig. 3A). However, these differences in overall plankton community were not the result of changes in diversity given that neither the Shannon’s Index of Diversity (p = 0.607) nor the Simpson’s Indexes of Diversity (p = 0.552) showed significant differences in plankton community diversity across time of day (Fig. 3B, C).

**Fig. 3.**
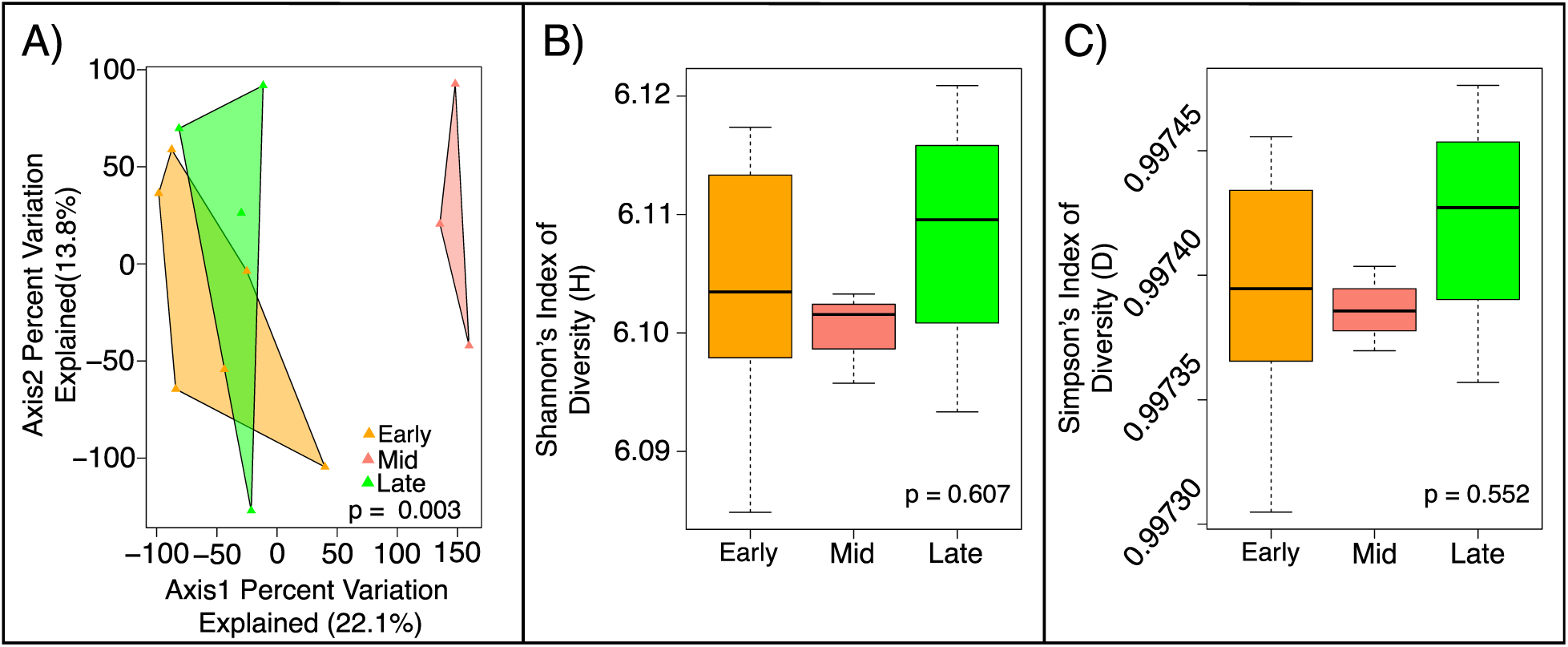
STRI Point plankton samples clustered by time of day (Early, Mid-day, Late) (A) Principal coordinate analysis of plankton communities by time of day at STRI Point. Percentages on each axis indicate the amount of variation explained by each axis. Early = orange, Mid-day = pink and Late = green. *Adonis* P-value demonstrates that there was a significant statistical difference in community composition across reef zones. (B) Mean Shannon and (C) Simpson diversity of plankton communities based on time of day. P-values demonstrate that there were no statistical differences in diversity across time of day and error bars represent the minimum and maximum indices of diversity.

Heatmaps of the most differentially abundant ASVs highlight the taxonomic orders driving the observed overall community shifts between sampling timepoints (Fig 4A, B). We observe several phytoplankton ASVs (Orders Bacillariophytina [diatoms], Syndiniales, and Peridiniphycidae) that are enriched mid-day compared to early or late, while other photosynthetic taxa (e.g., other Bacillariophytina and Zoantharia [zoanthids]) are enriched in the early and late timepoints, highlighting the complexity of diel vertical migrations observed on these reefs.

**Fig. 4.**
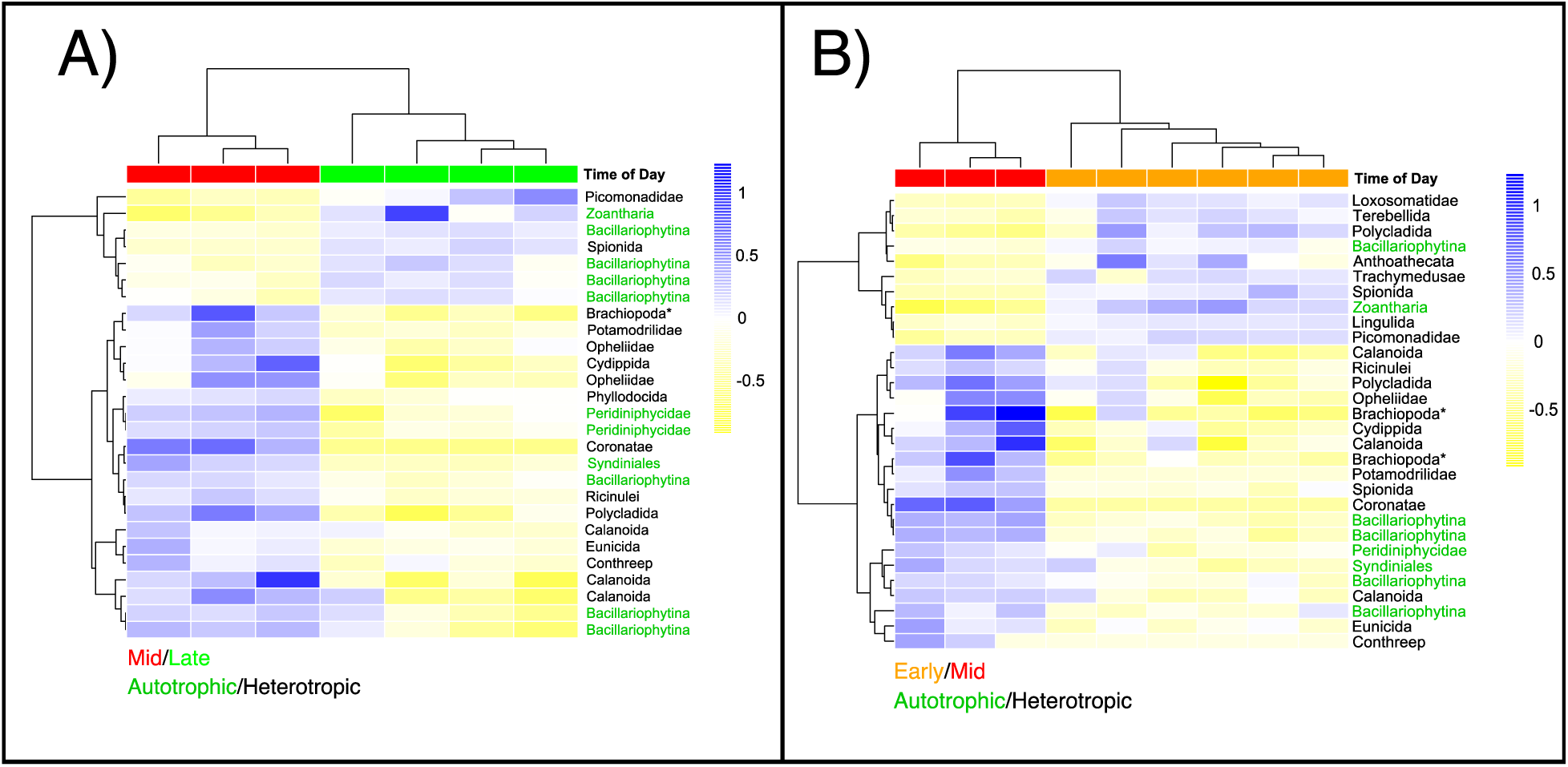
Diel Variation in specific plankton taxa. (A–B) Heatmaps of the most differentially abundant taxa between early and mid-day sampling. Asterisks are for ASVs that could only be identified to its phylum level instead of order like the rest. (A) and between mid-day and late sampling (B). Red, orange and green blocks indicate that libraries originated from plankton communities collected mid-day, early or late, respectively. Columns represent unique plankton tows and rows represent differentially abundant taxa. Taxa listed in black are heterotrophic whereas taxa listed in green are autotrophic. The color scale is in log2 (blue: enriched, yellow: depleted) and taxa and samples are clustered hierarchically based on Pearson’s correlation of their relative abundance across samples.

## DISCUSSION

### Environmental Differences Across Reef Zones

It has been well established that inshore and offshore reef zones differ in their environmental conditions across space and time. Inshore reefs experience increased turbidity, sedimentation, nutrients, and temperature variation, while offshore reefs are characterized by more moderate temperatures and lower turbidity as they are buffered by the open ocean (Boyer & Briceño 2011; Lirman & Fong 2007; Lirman et al. 2011). Here we show that the inshore and offshore reef zones on coral reefs in the Bocas del Toro Archipelago, Panamá are consistent with these expected environmental differences, with inshore reefs exhibiting lower light levels and warmer temperatures compared to offshore reef sites (Fig. 1). Warmer inshore waters and higher turbidity (i.e. reduced light) are also consistent with *in situ* data measured on other Caribbean reef tracts, including Belize (Castillo et al., 2012; Baumann et al., 2016) and Florida (Kenkel et al., 2017; Rippe et al., 2018). These differences in mean mid-day light values and temperature are expected to drive niche specialization across marine environments, with specific taxa exhibiting preferences for distinct reef environments (Edwards et al. 2016; Andersson et al. 1994; Takasuka et al. 2005).

### Plankton Community Show Few Differences Across Reef Zones

Although we observed significant differences in light and temperature across inshore and offshore reef zones (Fig. 1), these environmental variations did not correspond with overall differences in plankton communities (Fig. 2). This result is surprising given that on larger scales, it has been estimated that variation in sea-surface temperature explains roughly 90% of the geographic variation in plankton diversity throughout the Atlantic Ocean (Rutherford et al. 1999) and it has been shown that plankton communities can be affected by even finer-scale environmental variations including depth, temperature and trophic state of the water (i.e. particulate concentration, nutrients, and chlorophyll-*a*) (Owen 1989). This lack of plankton community structure observed here may suggest that the processes structuring plankton communities in this archipelago operate at much larger spatiotemporal scales than the scale investigated here. However, there are also several confounding hypotheses that may serve to reconcile these results.

First, the results presented here are based solely on sequencing data from three single point measurements in space and time on a single day at a specific reef site. With the exception of STRI Point, we do not consider temporal changes at these sites across days, and it is possible that our point collections do not accurately represent the average plankton community observed at these sites. Given that previous work has shown that kinetic properties of water influence planktonic organization (Mackenzie and Leggett 1991) and marine plankton communities can be more dispersed in high-energy, turbulent environments (Haury et al. 1990), it is possible that weather related influences during the days of sampling (e.g., wind) could have acted to homogenize plankton communities across sites. It is also possible that samples taken in a different season, as in the study by (Huang et al. 2004) could yield different results. Furthermore, our collections were also conducted using a 60 µm net, which excludes the sampling of smaller organisms, so it is also possible that community differences exist at smaller size fractions that were outside of the scope of this study. Lastly, we only measured temperature and light to assess environmental differences across reef zones and it could be that other physical and biochemical properties that were not measured here are stronger drivers of these tropical coastal plankton communities, like nutrient runoff from the mainland (D’Croz et al. 2005) and long term climate change (De Stasio et al. 1996).

Another important consideration is that sequencing plankton communities introduces its own set of caveats, including the fact that rDNA copy number per cell varies by orders of magnitude across unicellular eukaryotes (e.g., dinoflagellates and ciliates) (Weider et al. 2005; Gong et al. 2013). Therefore, caution must be exercised when interpreting organism abundance based on rDNA sequence abundance. Variation in rDNA copy number can even occur within a species and a recent single-cell sequencing study found that rDNA and rRNA copy number scaled with cell size in two ciliate species (Fu and Gong 2017), so variation in plankton size, which was not measured here, could have influenced relative abundances. Equally plausible, plankton communities across these sites may be homogenous but the functional processes within each group of taxa may differ transcriptionally across sites, which has been previously observed in diatoms in response to iron availability (Cohen et al. 2017) and in dinoflagellates in response to light environment (Davies et al. 2018). We also leveraged 18S rDNA sequencing, which is known to be highly conserved across taxa, but this single locus approach overlooks any sort of within-species population genetic differences that may exist between sites (Rodríguez et al. 2005; Martiny et al. 2009).

Given these caveats, we propose that future studies should couple more traditional microscopy techniques with 18S rDNA sequencing and perhaps consider a multidisciplinary approach incorporating metatranscriptomics or population genetics of specific taxa of interest in order to capture potential ecological and functional differences between plankton communities across reef zones.

### Time of Day Played a Role in Structuring Plankton Communities

Despite not finding differences in overall plankton community structure across reef zones, we did observe differences across time of day within the STRI Point site. We observed differences in overall community structure (Fig. 3A), reduced variation in diversity indices (Fig. 3B,C), and differentially enriched taxa over the course of the sampling day (Fig. 4A,B), which are all likely the result of diel vertical migration (DVM) of both phytoplankton and zooplankton. DVM is the movement of plankton and fish vertically in the water column over a daily cycle. For zooplankton, these movements are most commonly (but not always) up to the surface at dusk and back to the deeper waters at dawn (Lampert 1989; Ohman 1990; Brierley 2014) in order to avoid predation pressures (Ohman 1988; Lampert 1989). Planktivorous fishes are visual hunters, and most species inhabiting nearshore environments have been found to feed during the day, thus exerting a diurnal predation pressure on plankton (Morgan 1990; Motro et al. 2005). Predation pressure of planktivorous fishes on zooplankton is also strong on coral reefs (Hamner et al. 1988), and has been shown to drive vertical patterns of zooplankton in these habitats (Motro et al. 2005). Specifically in Bocas del Toro, Kerr et al. (2014) demonstrated that predation risk is higher during the day than at night for *Artemia franciscana* nauplii. However, the temporal gradient in planktonic predation risk was dependent on prey life history stage (i.e., size), as adult *A. franciscana* did not show predation differences across the diurnal cycle (Kerr et al. 2014). While the “normal” zooplankton migration is considered to be ascending in the evening and descending in the morning, examples of “reversed” migrations are also common (Lampert 1989; Ohman 1990), with migration patterns varying by whether predation pressure is from visually hunting planktivorous fishes or nocturnally feeding zooplankton (Ohman 1990). Here, we find evidence for both normal and reversed patterns of zooplankton migration at the STRI Point site, with some zooplankton taxa enriched at mid-day and some enriched earlier/later in the day.

We also found evidence of phytoplankton DVM in Bocas del Toro, as some taxa were enriched at mid-day and some were enriched in either the morning or evening. Phytoplankton DVM is generally understood as a mechanism for these organisms to optimize light and nutrient gradients, therefore moving into shallower waters during the day to photosynthesize and moving deeper in the water column at night to uptake nutrients (Raven and Richardson 1984; Ault 2000). As photosynthetic characteristics of phytoplankton vary, optimum depth and migration pattern varies by the underwater light pattern and the organism being considered (Ault 2000). As with zooplankton, we found evidence of both the normal migration and reversed migration patterns. Some phytoplankton sampled appeared to be migrating up to surface waters at mid-day (e.g., Bacillariophytina [diatoms], Syndiniales, and Peridiniphycidae). However, other taxa (e.g., other Bacillariophytina and Zoantharia [zoanthids]) were enriched in the morning/evening, suggesting that they were migrating away from surface waters during mid-day (Fig 5). This reversed pattern is likely evidence of these organisms migrating away from high noon-time irradiance in order to avoid photoinhibition (Anderson and Stolzenbach 1985; Kingston 1999; Flynn and Fasham 2002). However, as our analyses can only distinguish to the level of Order, it is difficult to interpret the factors influencing the patterns observed for both phytoplankton and zooplankton.

**Fig. 5.**
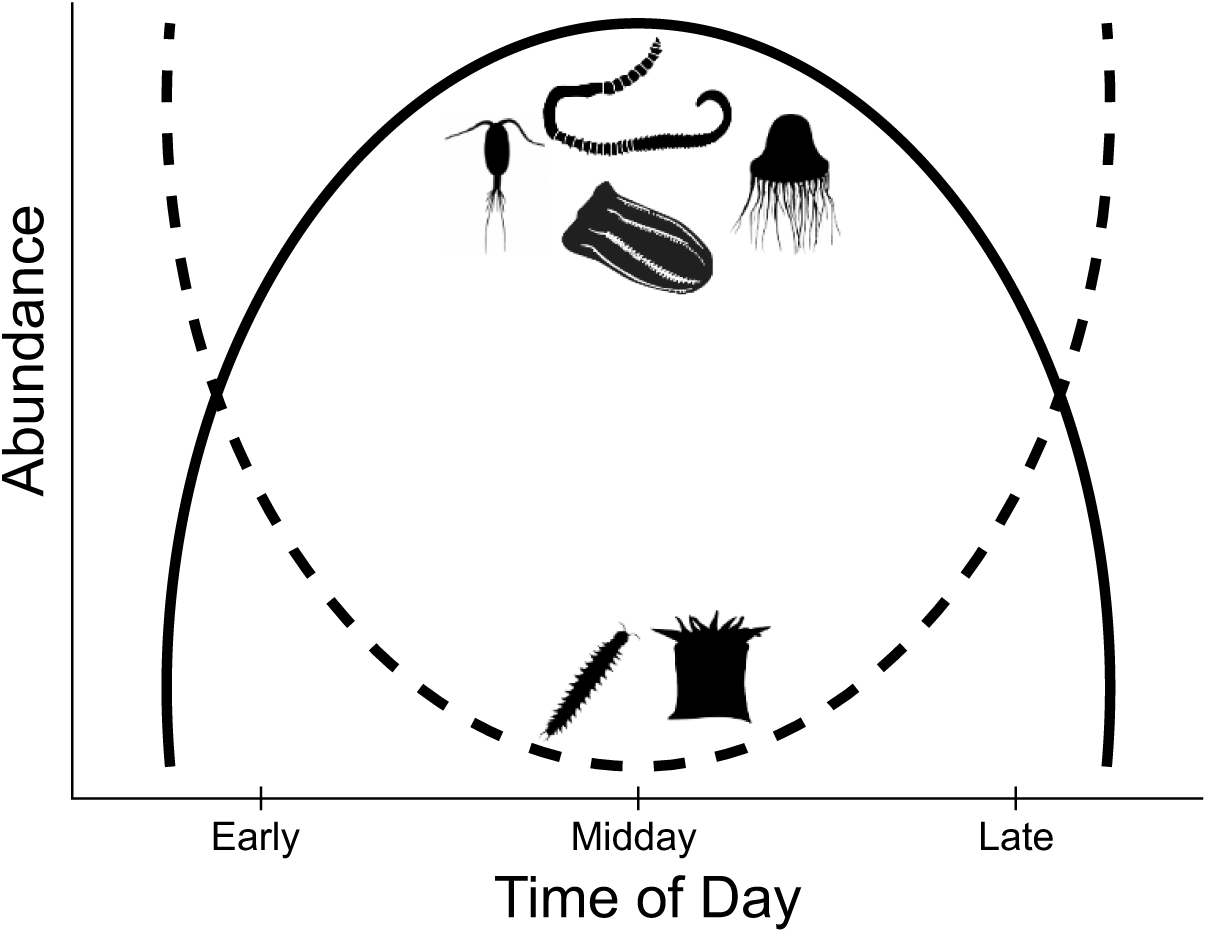
Conceptual Model for Diel Community Shifts in Plankton Communities at STRI Point. Based on our data, Cydippida, Eunicida, Coronatae, and Calanoida taxa are more abundant mid-day, which are represented by the solid line, and Spionida and Zoantharia are more abundant during early and late time frames, which are represented by the dashed line.

### Concluding Thoughts

Tracking plankton community composition through space and time is critical as climate change progresses. Correlating the environmental conditions experienced in Bocas del Toro to the plankton communities across these sites builds a baseline dataset upon which future studies can build in order to assess how changing environments are influencing these communities. Current estimates suggest that the oceans have warmed by ca. 0.6°C over the past 100 years (IPCC, 2007) and have absorbed almost 50% of all the anthropogenic CO2 emitted over the last 250 years (Sabine *et al*., 2004), and as the oceans continue to change, the need to characterize baseline community structure is imminent. Plankton not only play a central and critical role in the health and productivity of the oceans, but can also serve as sensitive indicators of climate change. As plankton communities shift in response to climate change, the availability of energy for other trophic levels will also shift, which will undoubtedly modulate food web dynamics. Therefore, a more comprehensive description of baseline plankton communities provided by studies like the one presented here are needed before we can make accurate projections of what impacts these climate-mediated shifts in plankton communities will have on future reefs.

## Supporting information

ESM

## Conflict of Interest

On behalf of all authors, the corresponding authors state that there are no conflicts of interest.

## Acknowledgements

We acknowledge the Government of Panamá, Ministerio de Ambiente and STRI for all permitting (#SE/A-28-15 and #SEX/AO-2-15) and coordination of fieldwork. We thank the Marchetti Laboratory at the UNC Chapel Hill for providing molecular lab space and thoughtful discussions. Lastly, we thank J.P Rippe, Colleen Bove, Justin Baumann, and Clare Fieseler for fieldwork assistance. Funding was provided by the National Science Foundation (NSF) grant OCE-1459522 to K.D.C, a UNC Summer Undergraduate Research Fellowship to L.K.B., A.M.R. was supported by NSF-REU grant BIO-1659605, and start-up funds from Boston University to S.W.D.

