## Supplementary material for "Eukaryotic plankton community stability across reef environments in Bocas del Toro Archipelago (Panamá)": ESM

Electronic Supplementary Material

**ESM Figure 1**:

Graphical representation of the library preparation and downstream analysis steps utilized in this study. Original primers (P.) and linkers were used for amplification of 18s rDNA region for compatibility with Illumina MiSeq. The second PCR had the addition of unique barcodes (BC) and Illumina adapters (P5) for species identification. *DADA2* analysis was used to analyze data and these results were compared to the SILVA database in order to identify the specific taxa in the samples.

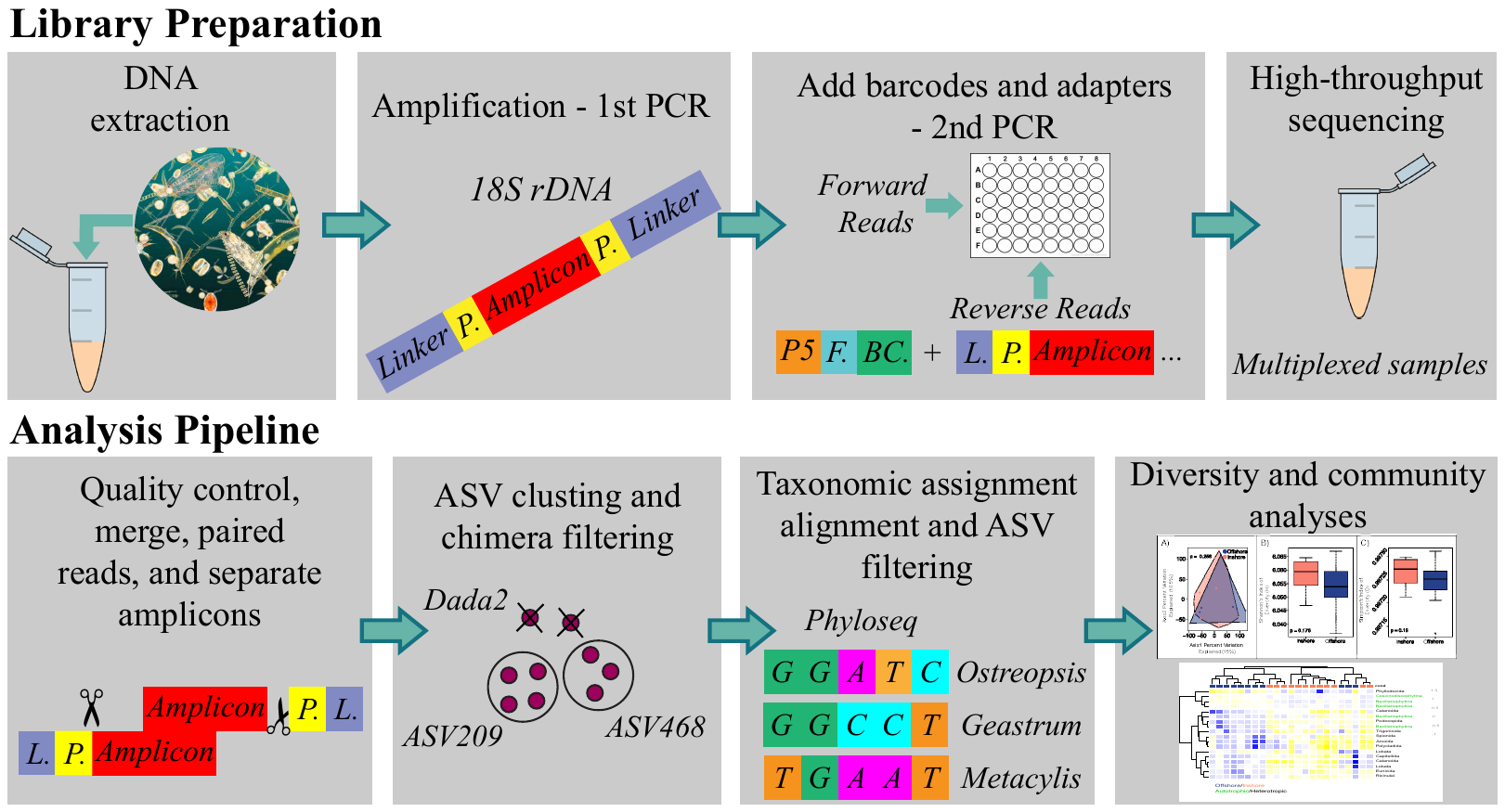

**ESM Figure 2**:

Variation in plankton samples across sites collected mid-day**.** (A) Principal coordinate analysis of plankton communities by individual sites. Percentages on each axis indicate the amount of variation explained by each axis. Inshore sites = salmon, Offshore sites = blue. *Adonis* P-value demonstrates that there was no significant statistical difference in community composition across reef sites. (B) Mean Shannon and (C) Simpson diversity of the plankton communities across individual sites. P-values demonstrate that there were no statistical differences in diversity across sites and error bars represent the minimum and maximum indices of diversity.

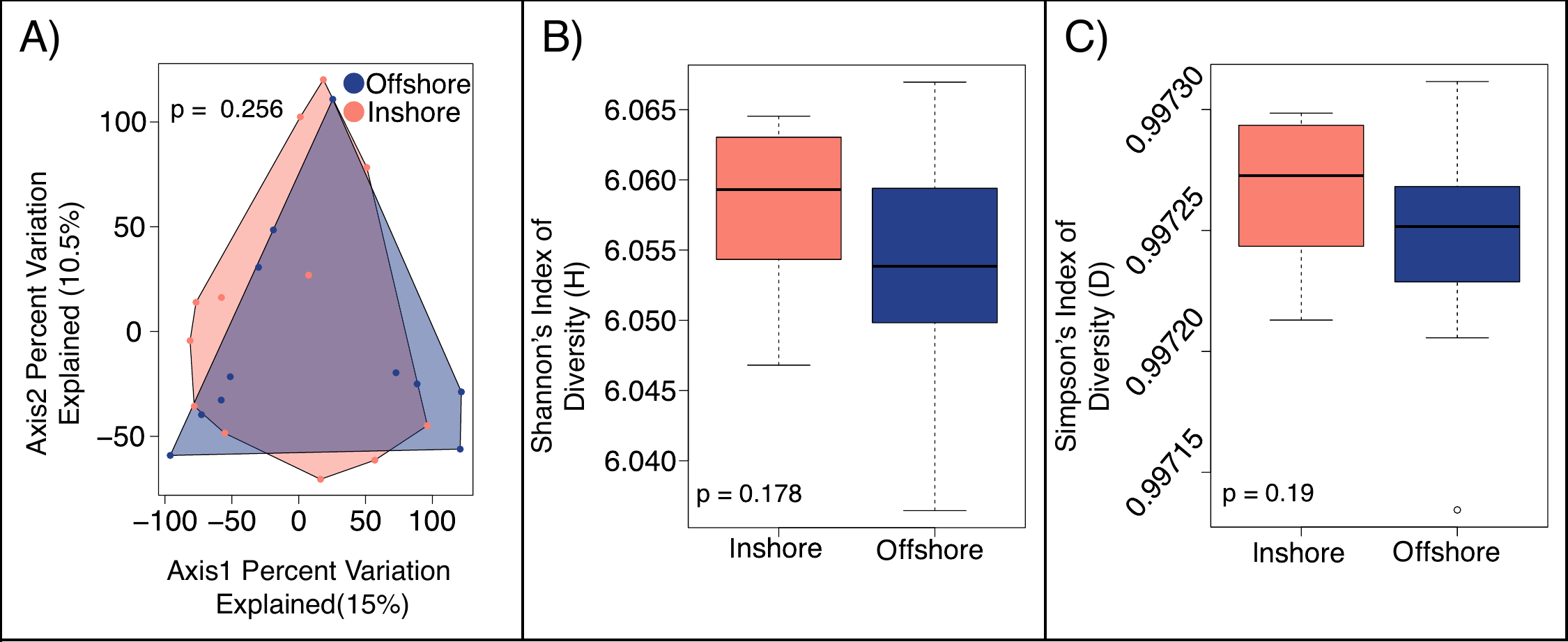

**ESM Table 1**:

Supplemental Table S1. Metadata for individual library preparations including the sample ID, site of collection, time of collection, number of PCR cycles used to amplify the sample, raw read count and the number of reads resulting from each step of the *Dada2* analysis pipeline with final efficiency values as a percentage of the original reads.

| **Sample ID** | **Site Name** | **Time** | **Number of Cycles** | **Raw Reads** | **Filtered** | **Merged** | **Tabled** | **No Chim** | **Mapping Efficiency** |
| --- | --- | --- | --- | --- | --- | --- | --- | --- | --- |
| P28-1 | Cristobal | Mid | 23 | 196392 | 164632 | 80832 | 80780 | 80532 | 41% |
| P28-2 | Cristobal | Mid | 26 | 84457 | 70369 | 34062 | 34051 | 33871 | 40% |
| P29-1 | Cristobal | Mid | 23 | 124361 | 103493 | 48901 | 48844 | 48647 | 39% |
| P29-2 | Cristobal | Mid | 25 | 114043 | 97011 | 45575 | 45550 | 45393 | 40% |
| P30-1 | Cristobal | Mid | 23 | 155171 | 115284 | 56913 | 56880 | 56679 | 37% |
| P30-2 | Cristobal | Mid | 25 | 137683 | 115632 | 60446 | 60404 | 60220 | 44% |
| P16-1 | Punta Laurel | Mid | 25 | 150442 | 94131 | 43651 | 43641 | 43474 | 29% |
| P16-2 | Punta Laurel | Mid | 29 | 83571 | 52128 | 25037 | 25021 | 24886 | 30% |
| P17-1 | Punta Laurel | Mid | 25 | 160054 | 134464 | 66339 | 66277 | 66093 | 41% |
| P17-2 | Punta Laurel | Mid | 29 | 159768 | 134403 | 67427 | 67351 | 67127 | 42% |
| P18-1 | Punta Laurel | Mid | 25 | 168019 | 140098 | 70338 | 70293 | 70098 | 42% |
| P18-2 | Punta Laurel | Mid | 25 | 144179 | 119403 | 57813 | 57780 | 57560 | 40% |
| P34-1 | Punta Donato | Mid | 24 | 72853 | 55894 | 27175 | 27143 | 26969 | 37% |
| P34-2 | Punta Donato | Mid | 28 | 81541 | 68269 | 37713 | 37694 | 37556 | 46% |
| P35-1 | Punta Donato | Mid | 23 | 158414 | 132451 | 65306 | 65260 | 65051 | 41% |
| P35-2 | Punta Donato | Mid | 23 | 112475 | 95638 | 48650 | 48621 | 48474 | 43% |
| P36-1 | Punta Donato | Mid | 24 | 170227 | 142018 | 69281 | 69201 | 69031 | 41% |
| P36-2 | Punta Donato | Mid | 29 | 112689 | 94452 | 43990 | 43942 | 43736 | 39% |
| P19-1 | STRI Point | Early | 25 | 203228 | 172445 | 85613 | 85465 | 85205 | 42% |
| P19-2 | STRI Point | Early | 26 | 163277 | 137402 | 66280 | 66175 | 65962 | 40% |
| P20-1 | STRI Point | Early | 25 | 149034 | 124438 | 62666 | 62622 | 62406 | 42% |
| P20-2 | STRI Point | Early | 26 | 77653 | 64941 | 33048 | 33015 | 32849 | 42% |
| P21-1 | STRI Point | Early | 23 | 250787 | 203841 | 102260 | 102188 | 101879 | 41% |
| P21-2 | STRI Point | Early | 26 | 105611 | 81089 | 42188 | 42172 | 42018 | 40% |
| P25-1 | STRI Point | Early | 24 | 151846 | 123721 | 67265 | 67206 | 66977 | 44% |
| P25-2 | STRI Point | Early | 26 | 91175 | 76330 | 42510 | 42476 | 42312 | 46% |
| P26-1 | STRI Point | Early | 26 | 197363 | 167535 | 80343 | 80243 | 79987 | 41% |
| P26-2 | STRI Point | Early | 26 | 191465 | 148694 | 77023 | 76977 | 76752 | 40% |
| P27-1 | STRI Point | Early | 24 | 138317 | 94551 | 47140 | 47121 | 46978 | 34% |
| P27-2 | STRI Point | Early | 26 | 143583 | 99166 | 50232 | 50199 | 50012 | 35% |
| P37-1 | STRI Point | Mid | 23 | 132738 | 110027 | 52161 | 52111 | 51920 | 39% |
| P37-2 | STRI Point | Mid | 27 | 143909 | 111289 | 52790 | 52769 | 52595 | 37% |
| P38-1 | STRI Point | Mid | 23 | 182137 | 154148 | 71435 | 71371 | 71155 | 39% |
| P38-2 | STRI Point | Mid | 27 | 126716 | 81252 | 36090 | 36081 | 35915 | 28% |
| P39-1 | STRI Point | Mid | 23 | 134975 | 110718 | 52029 | 51980 | 51794 | 38% |
| P39-2 | STRI Point | Mid | 27 | 128897 | 108186 | 50295 | 50260 | 50093 | 39% |
| P22-1 | STRI Point | Late | 21 | 72103 | 55000 | 28756 | 28732 | 28580 | 40% |
| P22-2 | STRI Point | Late | 26 | 99875 | 66386 | 35070 | 35062 | 34881 | 35% |
| P23-1 | STRI Point | Late | 25 | 177228 | 148186 | 74929 | 74881 | 74659 | 42% |
| P23-2 | STRI Point | Late | 26 | 103218 | 87518 | 44749 | 44712 | 44566 | 43% |
| P24-1 | STRI Point | Late | 24 | 184760 | 154540 | 78127 | 78086 | 77892 | 42% |
| P24-2 | STRI Point | Late | 26 | 119937 | 102506 | 52845 | 52812 | 52651 | 44% |
| P31-1 | STRI Point | Late | 21 | 140949 | 91237 | 44612 | 44584 | 44390 | 31% |
| P31-2 | STRI Point | Late | 25 | 49185 | 38466 | 22160 | 22152 | 21968 | 45% |
| P32-1 | STRI Point | Late | 23 | 223952 | 185843 | 89631 | 89554 | 89264 | 40% |
| P32-2 | STRI Point | Late | 27 | 100172 | 84280 | 43408 | 43392 | 43259 | 43% |
| P33-1 | STRI Point | Late | 23 | 143196 | 119727 | 58978 | 58941 | 58777 | 41% |
| P33-2 | STRI Point | Late | 29 | 98528 | 82854 | 43399 | 43362 | 43203 | 44% |
| P04-1 | Bastimentos North | Mid | 25 | 94604 | 60848 | 27483 | 27454 | 27311 | 29% |
| P04-2 | Bastimentos North | Mid | 29 | 172685 | 147336 | 65893 | 65818 | 65604 | 38% |
| P05-1 | Bastimentos North | Mid | 25 | 104093 | 87668 | 40505 | 40485 | 40365 | 39% |
| P05-2 | Bastimentos North | Mid | 29 | 124644 | 103230 | 46842 | 46802 | 46657 | 37% |
| P06-1 | Bastimentos North | Mid | 25 | 129057 | 109805 | 47532 | 47479 | 47327 | 37% |
| P06-2 | Bastimentos North | Mid | 29 | 186888 | 156681 | 70847 | 70806 | 70589 | 38% |
| P07-1 | Bastimentos South | Mid | 26 | 85891 | 59240 | 27253 | 27233 | 27059 | 32% |
| P07-2 | Bastimentos South | Mid | 29 | 57896 | 45106 | 20413 | 20401 | 20194 | 35% |
| P08-1 | Bastimentos South | Mid | 26 | 139999 | 116589 | 55043 | 55001 | 54830 | 39% |
| P08-2 | Bastimentos South | Mid | 29 | 133845 | 112875 | 51632 | 51598 | 51412 | 38% |
| P09-1 | Bastimentos South | Mid | 25 | 48267 | 36802 | 16188 | 16158 | 16007 | 33% |
| P09-2 | Bastimentos South | Mid | 29 | 133589 | 111798 | 52044 | 52024 | 51849 | 39% |
| P13-1 | Drago Mar | Mid | 24 | 108825 | 90680 | 43795 | 43767 | 43605 | 40% |
| P13-2 | Drago Mar | Mid | 29 | 135827 | 114345 | 53477 | 53442 | 53250 | 39% |
| P14-1 | Drago Mar | Mid | 25 | 131828 | 110115 | 50233 | 50130 | 49968 | 38% |
| P14-2 | Drago Mar | Mid | 29 | 74162 | 62263 | 29686 | 29648 | 29501 | 40% |
| P15-1 | Drago Mar | Mid | 25 | 162091 | 118777 | 56658 | 56609 | 56434 | 35% |
| P15-2 | Drago Mar | Mid | 29 | 94410 | 72386 | 35543 | 35525 | 35395 | 37% |
| P10-1 | Popa Island | Mid | 24 | 99012 | 81388 | 37579 | 37561 | 37407 | 38% |
| P10-2 | Popa Island | Mid | 29 | 119462 | 96483 | 45109 | 45087 | 44927 | 38% |
| P11-1 | Popa Island | Mid | 20 | 102479 | 85662 | 39829 | 39802 | 39639 | 39% |
| P11-2 | Popa Island | Mid | 23 | 152471 | 126713 | 59297 | 59254 | 59057 | 39% |
| P12-1 | Popa Island | Mid | 20 | 87374 | 72338 | 34294 | 34280 | 34126 | 39% |
| P12-2 | Popa Island | Mid | 23 | 165451 | 138884 | 64649 | 64598 | 64352 | 39% |
