## Supplementary figures and images for "Eukaryotic plankton community stability across reef environments in Bocas del Toro Archipelago (Panamá)"

### FigS1

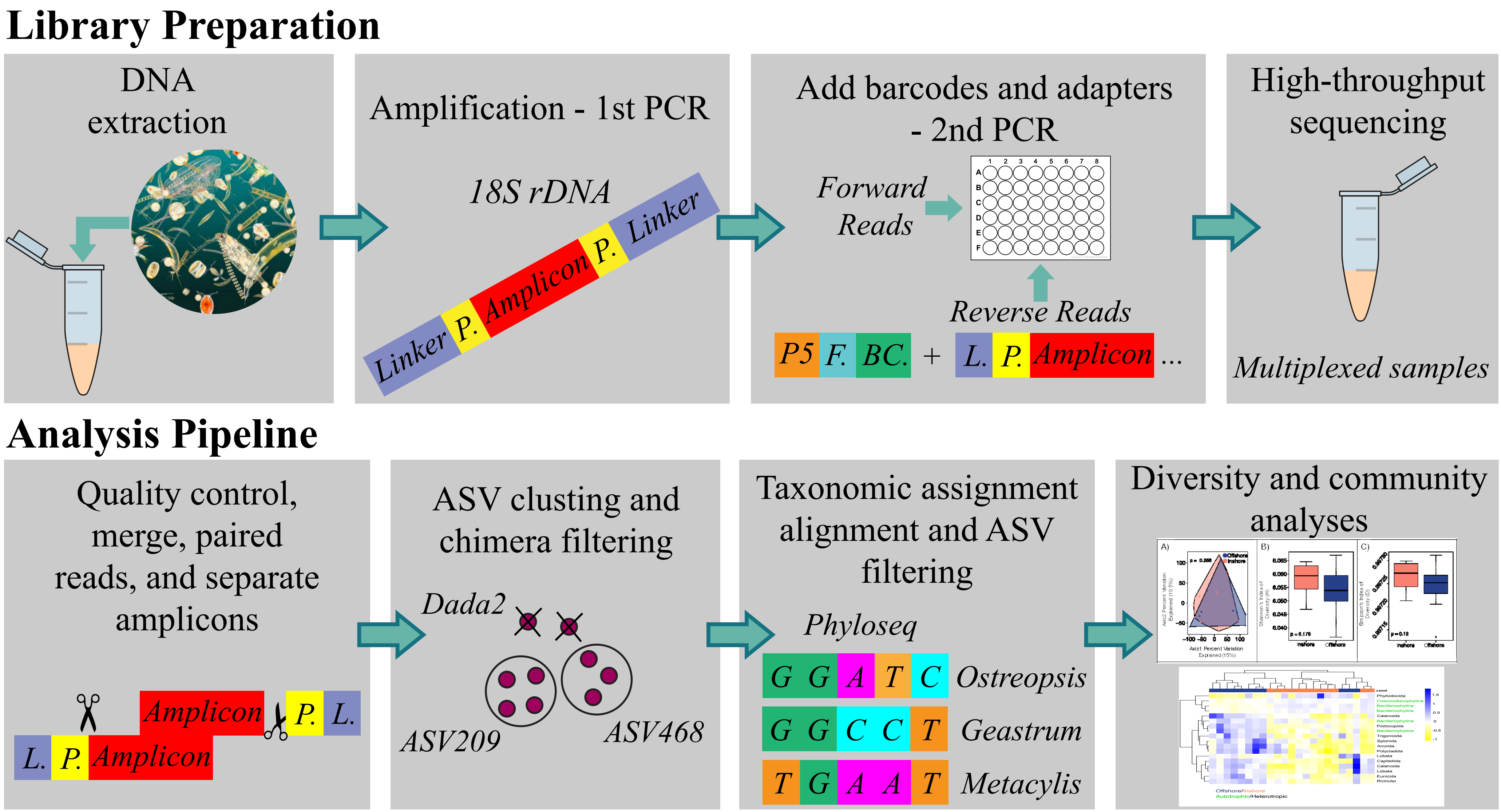

### FigS2

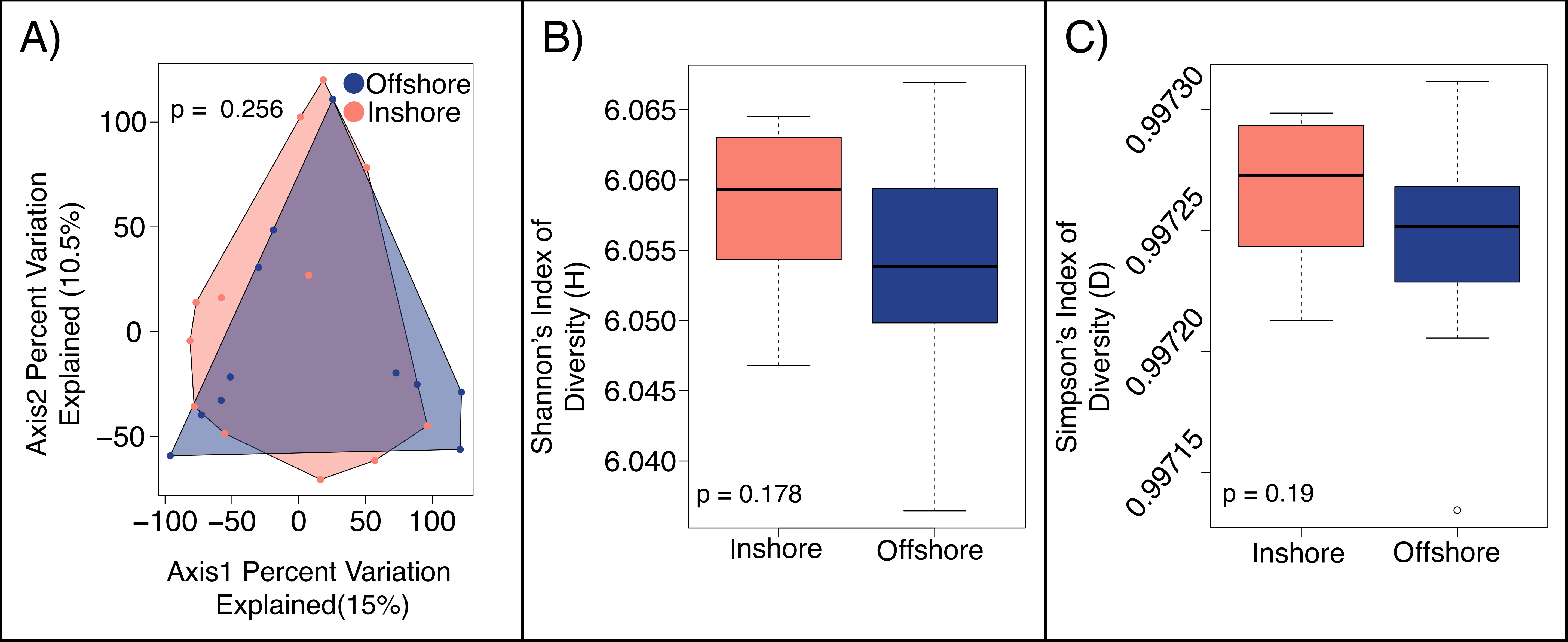
